# Trophotypes of the human gut microbiome: discrete energetic states under a macroecological framework

**DOI:** 10.64898/2026.05.27.728235

**Authors:** Manuel Mendoza

## Abstract

A central prediction of complex-systems ecology is that strongly interacting communities settle into a limited number of recurrent configurations — attractors of their internal energy-flow dynamics — rather than spreading continuously across the space of possible compositions. This has been confirmed in terrestrial mammals, in birds and mammals combined at global scale, and in marine communities, using guild richness as a state descriptor that integrates long-term energetic capacity. Here I extend the framework to the human gut microbiome. Using 8,960 faecal samples from the American Gut Project described by the richness of twelve metabolic guilds, I apply Average Membership Degree analysis (AMD) and identify four discrete Trophotypes separated by sparsely populated regions of the functional space. Principal component analysis, applied independently to the same matrix, identified the same three guilds --- primary generalist degraders, butyrate producers, and acetate producers --- as the main axes of variation, together accounting for 78% of total variance; a diagnostic random forest recovered the same partition structure. The four Trophotypes occupy the four quadrants of the energetic plane defined by input through primary degradation and retention through butyrate production, in a topology consistent with multiple stable configurations sustained by stoichiometric constraints. Host-level metadata predict membership weakly across three independent algorithms (Cohen’s κ between 0.09 and 0.13). The human gut microbiome organises into a small set of recurrent functional states aligned with broad energetic axes, consistent with the multistability expected in systems governed by nonlinear network interactions.

**Importance:** The human gut microbiome does not exist in a single state: different individuals harbour communities with qualitatively different functional organisations. Understanding why requires knowing how many distinct configurations exist and what governs them. This study applies a geometric framework previously used to characterise trophic structures of bird and mammal communities at global scale to a large cohort of human faecal microbiomes. Four discrete functional configurations emerge, defined by variation in the capacity to input energy through primary polysaccharide degradation and to retain it through butyrate and acetate production — the two macroscopic axes that current theory identifies as the organising dimensions of the colonic fermentative network. Self-reported host metadata predict configuration membership only weakly, indicating either that the relevant control variables are not captured by survey data or that the configurations represent genuinely multistable attractors of the system.

## Introduction

Communities of strongly interacting species are dynamical systems whose state evolves under non-linear feedbacks among components, environmental constraints, and the flow of energy and matter across trophic levels. Theory predicts that such systems should not occupy the space of possible compositions uniformly; instead, they should settle into a limited number of recurrent configurations — attractors of the underlying dynamics — separated by sparsely populated transitional regions (1–3). This expectation has been a central tenet of complex-systems ecology for decades but has remained difficult to test empirically at the scale of whole communities, in part because energy flows across trophic guilds cannot be measured directly, especially at the spatial and taxonomic resolution required to detect the predicted structure.

A practical solution emerged from the recognition that the richness of each functional guild — the number of coexisting species or operational units capable of processing a given class of resources — can serve as a surrogate for the energy that the guild is able to process: higher resource availability supports larger viable populations and, indirectly, a larger number of coexisting species. Richness, unlike relative abundance, is comparatively insensitive to short-term fluctuations: it reflects the long-term capacity of the community to sustain a given metabolic process, a time scale closer to that of the dynamics shaping the underlying attractor than to its momentary occupation (4). The trophic structure of a community — defined by the vector of guild richness values — thus provides a tractable state descriptor in which the predicted discrete configurations can be detected directly.

Recent work has confirmed this prediction in vertebrate communities at broad biogeographical scales, using the Average Membership Degree (AMD) framework to identify recurrent configurations as regions of high local density in the guild-richness state space (5, 8). Six trophic structures emerged when AMD was applied to large mammals worldwide (5); the same six were recovered when the analysis was extended to all terrestrial mammals and to non-marine birds, separately and combined, despite substantial differences in species number and guild definitions (6). Across these systems, the six structures align with an increasing gradient of net primary productivity, identifying climate-mediated energy availability as the dominant axis along which trophic structures are organised. That the same pattern is recovered across taxonomic groups and guild definitions suggests that the discrete structure reflects general ecological processes rather than artefacts of a particular classification. The same framework applied to marine vertebrate communities at global scale recovered a small set of recurrent configurations (7), suggesting that the property generalises across terrestrial and aquatic systems.

If the property of organising into a small number of recurrent configurations is generic for complex communities with non-linear feedbacks, it should hold for microbial systems as well. The human gut microbiome is a particularly informative test case: hundreds of taxa interact through a dense web of cross-feeding, competitive and metabolic dependencies whose thermodynamic structure has been characterised in some detail (9), and theoretical models predict the existence of multiple stable configurations sustained by stoichiometric constraints on energy and matter flows (10). Empirically, discrete clusters in the composition of the human gut microbiome have been documented since the original description of enterotypes (11), although subsequent work showed that taxonomic discreteness is less sharp than initially proposed and depends substantially on methodological choices (12, 13). A possible reason is that the discreteness predicted by complex-systems theory operates on the functional organisation of the community — the network of energy flows through metabolic guilds — rather than on taxonomic identity per se, a contingent outcome of evolutionary history and dispersal. Taxonomic clusters, when observed, may correspond to partial projections of underlying functional configurations onto the taxonomic axis, mediated by the loose association between taxonomy and feeding strategy in microbial systems.

Here I apply the macroecological framework that has proved informative for vertebrate communities to the human gut microbiome, asking whether the system organises into a small number of discrete configurations in functional space, and whether those can be interpreted in terms of the energetic variables predicted by current theory of microbial community dynamics. I use 8,960 faecal samples from the American Gut Project (14), describing each by the richness of twelve metabolic guilds spanning the principal trophic roles in colonic fermentation. I apply AMD to detect recurrent configurations and a random forest to characterise the internal organisation of the partition. Principal component analysis provides an independent line of evidence on the geometry of variance. I then examine whether host-level metadata predict configuration membership, and use Dirichlet-multinomial mixture modelling as a methodological benchmark.

## Materials and Methods

### Data source and sample selection

I used the public release of the American Gut Project (AGP, Qiita study 10317) (14), a citizen-science cohort of self-collected faecal samples processed through a standardised 16S rRNA gene amplicon protocol targeting the V4 region. Sequences were classified against Greengenes 13.8 at 97% identity using SortMeRNA. The released BIOM table contains 15,158 faecal samples.

Sequencing depth varies substantially across samples, and richness-based descriptors are sensitive to depth heterogeneity. To eliminate this confound, I restricted the analysis to samples with at least 15,000 reads across all OTUs, retaining 8,960 samples (59% of the released cohort). All subsequent analyses operate on this filtered set.

### Trophic guild assignment

OTUs were mapped to metabolic guilds via a curated dictionary linking bacterial genera to functional roles in colonic fermentation. Twelve guilds were defined to span the principal energetic pathways of the system: primary degraders specialised on plant fibre (*Prevotella*); primary degraders with generalist polysaccharide preference (*Bacteroides*); mucin degraders (*Akkermansia*); lactate producers (*Bifidobacterium*, *Lactobacillus*); butyrate producers (*Faecalibacterium* sensu lato, including the recently described segregate species *F. duncaniae*, *F. hattorii* and *F. gallinarum* (33); *Roseburia*; *Coprococcus*); acetate producers (*Blautia*, classified separately given its role as an acetogen producing acetate via the Wood–Ljungdahl pathway and its abundance in the AGP cohort; while *B. hydrogenotrophica* is hydrogenotrophic, the most prevalent gut species — *B. luti* and *B. wexlerae* — lack the formate dehydrogenase and electron-bifurcating hydrogenase required for autotrophic growth on H2 + CO2, and use formate as intraspecific electron carrier instead (31, 32)); resistant-starch degraders (*Ruminococcus*); propionate producers (*Phascolarctobacterium*, *Veillonella*, *Dialister*; the first and third produce propionate primarily via succinate decarboxylation, an activity dependent on succinate generated by primary degraders, whereas *Veillonella* converts lactate to propionate); methanogenic H2 consumers (*Methanobrevibacter*); sulfate-reducing H2 consumers (*Desulfovibrio*); asaccharolytic H2 consumers (*Bilophila*; in the gut, this genus consumes H2 primarily as part of sulfite reduction during taurine and isethionate metabolism, producing hydrogen sulfide as the principal end-product); and proteolytic and opportunistic organisms (*Escherichia*, *Fusobacterium*). The dictionary covers eighteen genera selected for their high abundance, their well-documented metabolic role and their broad coverage of the principal energy-flow nodes in the human colon. OTUs assigned to genera outside this dictionary were grouped under an “Other or Unknown” category and excluded from the guild-richness matrix.

For each sample, the richness of every guild was computed as the number of distinct OTUs with non-zero counts assigned to that guild. The resulting matrix (8,960 × 12) is the state descriptor used in all downstream analyses.

### Detection of recurrent configurations: Average Membership Degree (AMD)

To detect recurrent configurations in the guild-richness space, I applied the Average Membership Degree (AMD) framework (5–8), a geometric, distribution-free approach that exploits a property predicted by complex-systems theory for strongly interacting communities: data points should accumulate in a limited number of dense regions of the state space separated by sparsely populated transitions. For a candidate *c*, fuzzy *c*-means clustering returns for each sample a vector of membership degrees, and the mean across samples of the maximum membership, AMDi(*c*), summarises how compactly the data partition into *c* groups. In a homogeneous cloud, AMDi(*c*) grows monotonically and slowly with *c*; when a true discrete structure of *c\** configurations exists, AMDi(*c*) rises sharply to a peak at *c* = *c\** and then drops. The optimal *c*_opt is identified at the local maximum.

To quantify the geometric definition of the detected configurations, I used the σ-equivalent metric. The method generates synthetic data with *c*_opt isotropic Gaussian clusters of standard deviation σ in a unit hypercube of the same dimensionality and sample size as the real data, and finds the σ that reproduces the observed AMDi(*c*_opt). σ-equivalent is expressed as a percentage of the hypercube side. I estimated it by bootstrap with 30 replicates of the full calibration, each with an independent random seed; the reported estimate is the median across replicates with a non-parametric 95% confidence interval (2.5%, 97.5% quantiles).

I computed the AMD curve for *c* = 2 to *c* = 14 with 50 iterations per value. The integer cluster labels returned by AMD are arbitrary across runs; I relabelled the configurations a posteriori by ascending mean total guild richness, so that T1 always denotes the lowest-richness configuration.

### Diagnostic analysis of guild contribution

To characterise which guilds participate in the emergence of the Trophotypes, I fitted a random forest classifier (16) on the AMD assignments using the twelve guild-richness variables as predictors, with 500 trees and default *mtry*. The model was used as a permutation-based diagnostic of variable contribution to the existing partition, not as a predictive tool. Variable importance was assessed via the mean decrease in classification accuracy on out-of-bag samples.

### Principal component analysis

I performed a principal component analysis on the guild-richness matrix, centring but not scaling the variables. Scaling to unit variance was avoided because differences in variance among guilds are biologically informative: a guild with high variance contributes more to community dynamics than one with low variance, and standardising would erase this distinction. Loadings and explained variance were obtained from *prcomp* in base R. Samples were projected onto the plane defined by the first two components and coloured by their AMD-derived Trophotype assignment.

### Host metadata as candidate control parameters

I tested whether host-level metadata predict Trophotype membership using three independent algorithms. From the metadata fields available in the AGP release, I selected forty candidate predictors grouped into three tiers reflecting prior biological expectation: Tier 1 (five variables: bowel-movement frequency and quality, antibiotic history, dietary pattern, types of plants consumed); Tier 2 (twenty-seven demographic, anthropometric, dietary-frequency, gastrointestinal and probiotic-use variables); and Tier 3 (eight lifestyle, geographic, travel and weight-change variables). Each variable was cleaned individually, harmonising encoding inconsistencies, converting ordinal Likert-style responses to ordered factors, and coercing physiologically implausible numeric values to missing. Full cleaning rules are documented in Supplementary Table S1.

To balance predictor coverage against sample loss under listwise deletion, I performed a greedy leave-one-out coverage analysis, dropping at each step the variable whose removal yielded the largest gain in complete-case sample size. The procedure identified country of residence as the only variable with catastrophic marginal cost; its exclusion left thirty-nine predictors with an analytic dataset of 4,260 complete-record samples. The Trophotype distribution in this subset differed from the full filtered cohort by at most 3.0 percentage points per class, indicating that the subsample is representative.

On this dataset I fitted three classifiers: a random forest with 1,000 trees and inverse-frequency case weights (24); an evolutionary classification tree (25) with maximum depth eight and complexity penalty α = 1, restricted to the top twelve random-forest predictors plus the five Tier 1 variables and evaluated under 5-fold stratified cross-validation; and a gradient-boosted classifier (26) with hyperparameters tuned over a 48-configuration grid via 5-fold cross-validation. The primary performance metric was Cohen’s κ to account for class imbalance. I also fitted the random forest on a subcohort of participants reporting no antibiotic use in the past year, as the closest available proxy for the system in or near its stable attractor regime.

### Dirichlet-multinomial mixture modelling

As a methodological benchmark, I fitted Dirichlet-multinomial mixture models (DMM) (15) to the same guild-richness matrix for K = 1 to K = 12 mixture components, with fit assessed via the Laplace approximation to the model evidence and the Akaike (AIC) and Bayesian (BIC) information criteria. DMM is a probabilistic, parametric counterpart to AMD: it asks whether the data can be represented as a finite mixture of Dirichlet-multinomial distributions and selects the optimal number of components by the elbow or absolute minimum of an information criterion. I report the full criterion trajectories across the tested range and compare them to the AMD curve obtained on the same data.

### Reproducibility and data availability

All analyses were performed in R 4.4.1 (22). Random seeds were set globally and before each stochastic step. Principal packages: AMDconfigurations (8) for configuration detection; randomForest (23) and ranger (24) for random forest classification; evtree (25) for evolutionary trees; xgboost (26) for gradient boosting; DirichletMultinomial (15) for DMM; mclust (29) for the Adjusted Rand Index; and ggplot2 (28) for figures. The complete analysis pipeline, metadata cleaning specification, guild-richness matrix, Trophotype labels and genus-to-guild dictionary are deposited at Zenodo (21). The raw 16S sequences and original metadata are available via Qiita (study 10317).

## Results

### Four discrete configurations in the guild-richness space

The AMDi curve computed on the 8,960 high-depth faecal samples rose monotonically from c = 2 to a peak at c = 4 (AMDi = 0.256) and dropped monotonically at higher values (Fig. 1A). The peak is well separated from the monotonic background expected under uniform dispersion in a hypercube of equivalent dimensionality, indicating that the data partition into four well-defined regions of the guild-richness space. The σ-equivalent at c = 4 was 13.24% (median across 30 bootstrap replicates; 95% CI [11.46%, 14.38%]; Fig. 1B), comparable to values reported for trophic structures in mammal and bird communities at global scale (6) and consistent with genuine recurrent attractors rather than artefactual partitioning.

**Figure 1.**
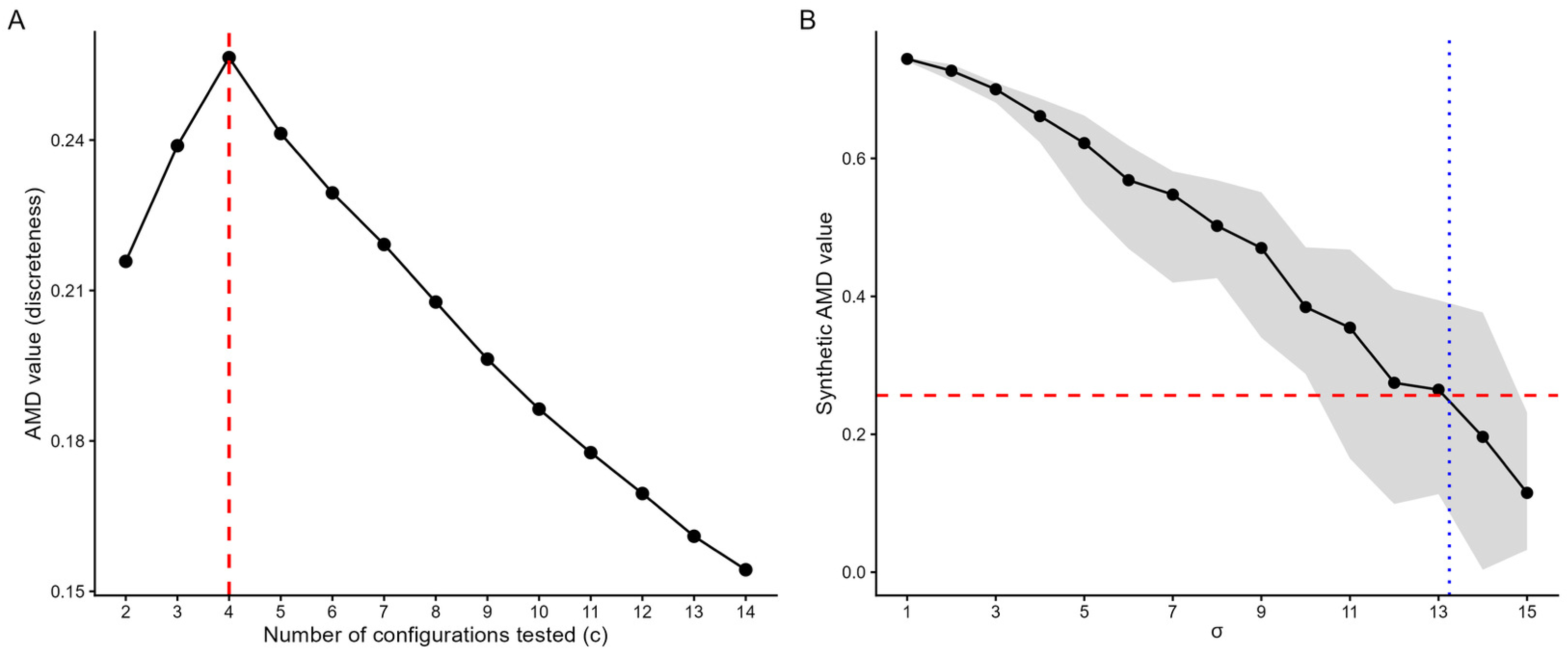
Detection of recurrent configurations in the guild-richness space and geometric calibration. (A) Average Membership Degree (AMDi) curve computed on the 8,960 high-depth faecal samples described by the richness of twelve metabolic guilds. AMDi(*c*) measures how compactly the data partition into *c* groups: the mean across samples of the maximum membership degree returned by fuzzy *c*-means clustering. The curve rises monotonically from c = 2 to a peak at c = 4 (AMDi = 0.256; red dashed line) and decreases monotonically thereafter, indicating that the data partition into four well-defined regions of the state space separated by sparsely populated transitional regions. (B) Bootstrap calibration of the σ-equivalent metric. For each value of σ on the grid, synthetic datasets containing four isotropic Gaussian clusters were generated 30 times with different seeds in a unit hypercube of the same dimensionality and sample size as the real data, and the AMDi peak at c = 4 was computed for each. The black curve shows the median synthetic AMDi across the 30 bootstrap replicates; the grey band shows the interquantile range between the 2.5% and 97.5% quantiles. The red dashed line marks the AMDi peak observed on the real data; the blue dotted line marks the median σ-equivalent (13.24% of the hypercube side length; 95% CI [11.46%, 14.38%]), the value of σ at which the synthetic calibration curve matches the real peak. The grey band remains bounded throughout the calibration range, indicating that the bootstrap estimate is robust to seed variability.

The four configurations, labelled T1 to T4 by ascending mean total guild richness, are populated as follows: T1, 2,164 samples (24.2%); T2, 2,461 (27.5%); T3, 2,346 (26.2%); T4, 1,989 (22.2%).

### Three guilds concentrate the organisation of the partition

Detection of the four Trophotypes by AMD does not by itself reveal how the discrete structure is organised internally. To identify which guilds participate in their emergence, I fitted a random forest classifier on the AMD assignments. The model was used as a diagnostic of variable contribution to the existing partition, not as a predictive tool — predicting AMD assignments from the variables that generated them would be tautological.

The variable-importance ranking is dominated by three guilds: primary generalist degraders (mean decrease in accuracy 368), butyrate producers (231) and acetate producers (144), with all other nine guilds yielding values below 50 (Fig. 2D). The drop from the third to the fourth predictor is roughly threefold, indicating that most of the discriminative information is concentrated in these three functional dimensions.

**Figure 2.**
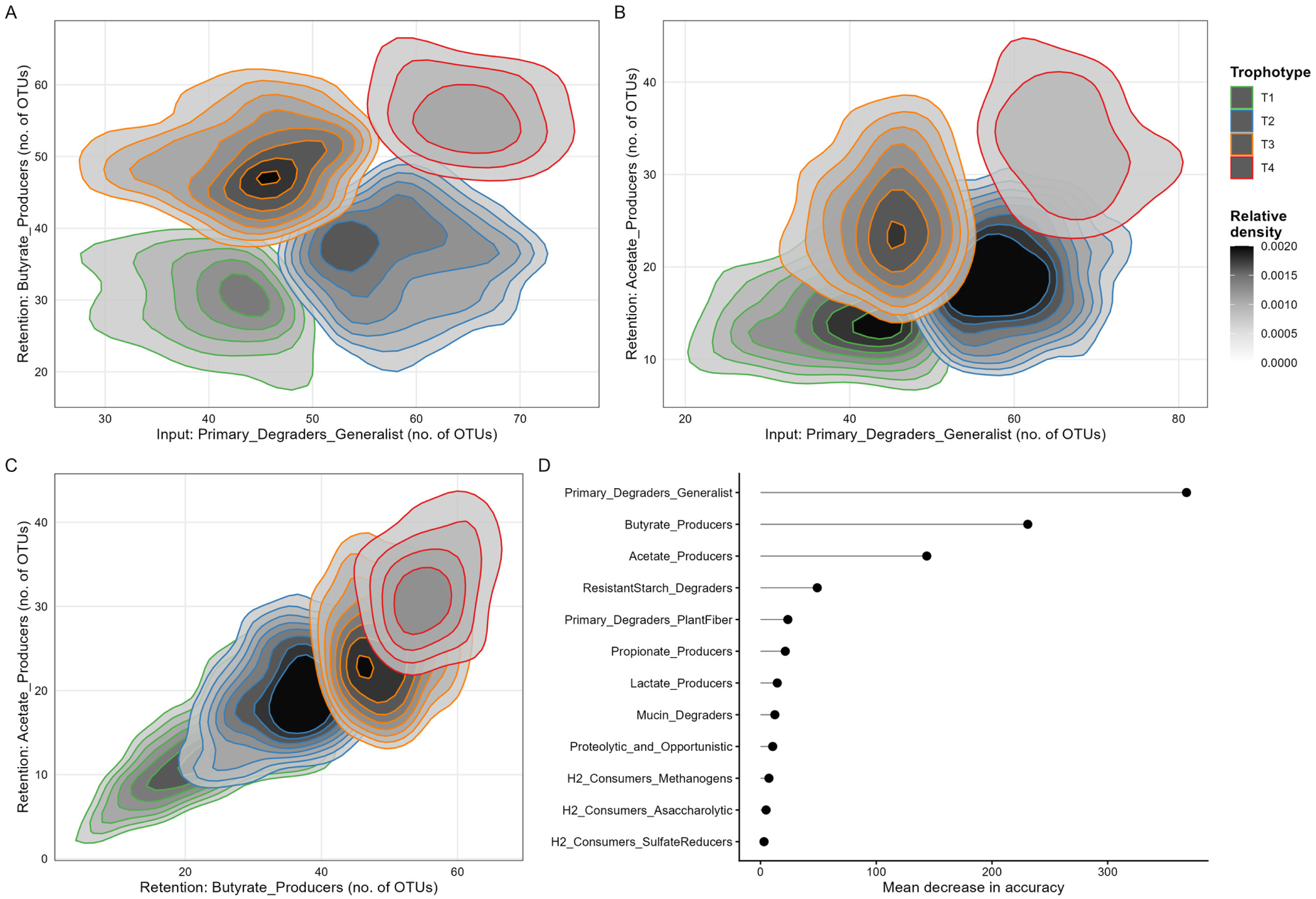
Functional structure of the Trophotypes. (A) Distribution of the 8,960 samples in the plane defined by primary generalist degraders (input of energy through polysaccharide degradation) on the *x*-axis and butyrate producers (retention of energy through butyrate production) on the *y*-axis. Each Trophotype is shown by density contours coloured according to the legend: T1 (green), T2 (blue), T3 (orange), T4 (red). The four configurations occupy the four quadrants of the energetic plane: T1 (low input, low retention), T2 (high input, low retention), T3 (low input, high retention) and T4 (high input, high retention). Grey shading indicates relative sample density. (B) Distribution of the same samples in the plane defined by primary generalist degraders on the *x*-axis and acetate producers (retention through the Wood–Ljungdahl pathway) on the *y*-axis. The four Trophotypes are distinguishable along both axes, with acetate separating T3 from T2 and isolating T4 from the remaining three configurations. (C) Distribution of the same samples in the plane defined by butyrate producers and acetate producers — the two retention guilds. The four configurations align along a positive diagonal, with the region of high acetate and low butyrate richness empty of samples. This geometric pattern is consistent with a thermodynamic coupling between butyrogenic fermentation and acetogenesis, although the precise mechanism — whether via direct H₂ transfer or via formate as electron carrier in the most prevalent gut Blautia species — remains under investigation. (D) Variable importance from a random forest classifier fitted on the AMD-derived Trophotype assignments using the twelve guild-richness variables as predictors. Importance is measured as the mean decrease in classification accuracy on out-of-bag samples under permutation of each variable. The model was used as a diagnostic of variable contribution to the existing partition, not as a predictive tool. Three guilds with direct energetic interpretation — primary generalist degraders, butyrate producers and acetate producers — concentrate the discriminative information, with all other guilds yielding values below 50.

In the plane defined by the two highest-importance guilds — primary generalist degraders (input through primary polysaccharide degradation) and butyrate producers (retention through short-chain fatty acid production) — the four Trophotypes occupy the four quadrants (Fig. 2A; Table 1). T1 sits at low values on both axes (low input, low retention); T2 combines high input with moderate retention; T3 shows the opposite (moderate input, high retention); T4 occupies the high-high region and is further distinguished by the highest mean acetate-producer richness in the cohort (38.5 vs 16.1 in T1). Input and retention are only partly coupled, giving rise to four qualitatively distinct combinations rather than a single ordered axis. The geometric basis of this organisation is examined in the next section.

**Table 1.** Mean guild richness per Trophotype. Mean number of distinct OTUs assigned to each metabolic guild, for each of the four Trophotypes identified by AMD on the high-depth faecal cohort (n = 8,960). Trophotypes are labelled T1 to T4 by ascending mean total guild richness. Values are rounded to one decimal place. Sample sizes per Trophotype: T1 = 2,164; T2 = 2,461; T3 = 2,346; T4 = 1,989. Guilds are ordered by mean richness in T4 (descending).

| Guild | T1 (n=2,164) | T2 (n=2,461) | T3 (n=2,346) | T4 (n=1,989) |
| --- | --- | --- | --- | --- |
| Primary_Degraders_Generalist | 32.8 | 62.0 | 39.9 | 71.1 |
| Butyrate_Producers | 25.5 | 33.7 | 50.8 | 60.2 |
| Acetate_Producers | 16.1 | 20.7 | 30.1 | 38.5 |
| ResistantStarch_Degraders | 10.6 | 13.2 | 16.3 | 21.0 |
| Primary_Degraders_PlantFiber | 10.1 | 8.0 | 8.9 | 11.3 |
| Propionate_Producers | 5.6 | 5.7 | 6.5 | 8.4 |
| Lactate_Producers | 5.5 | 5.0 | 6.8 | 7.7 |
| Mucin_Degraders | 1.6 | 1.8 | 1.6 | 2.1 |
| H<sub>2</sub>_Consumers_Asaccharolytic | 1.0 | 1.1 | 1.1 | 1.3 |
| H<sub>2</sub>_Consumers_SulfateReducers | 0.9 | 0.9 | 0.8 | 1.0 |
| Proteolytic_and_Opportunistic | 0.8 | 0.9 | 0.7 | 1.2 |
| H<sub>2</sub>_Consumers_Methanogens | 0.4 | 0.3 | 0.4 | 0.4 |

### The discriminative subspace is the principal subspace of the data

This concentration of discriminative information in three guilds could reflect either the main structure of variance in the data or simply a property of the AMD partition itself. To distinguish between the two, I performed PCA on the full twelve-dimensional matrix without reference to the AMD assignment.

PC1 and PC2 together explain 78.2% of the total variance (52.3% and 25.9%, respectively); PC3 adds 8.4%, dominated by an opposition between acetate and butyrate producers (loadings −0.83 and 0.54). PC1 is a positive combination of primary generalist degraders (0.66), butyrate producers (0.60) and acetate producers (0.40), with minor contributions from resistant-starch degraders (0.20) and all other guilds below 0.05 (Table S2); PC2 is dominated by primary generalist degraders (0.75) and butyrate producers (−0.53). The same three guilds highlighted by the random forest also dominate the empirical structure of variance. The projection of samples onto PC1 × PC2 coloured by AMD-derived Trophotype (Fig. 3) shows the four configurations as compact, contiguous regions of the principal plane, reproducing the four-quadrant arrangement of Fig. 2A.

**Figure 3.**
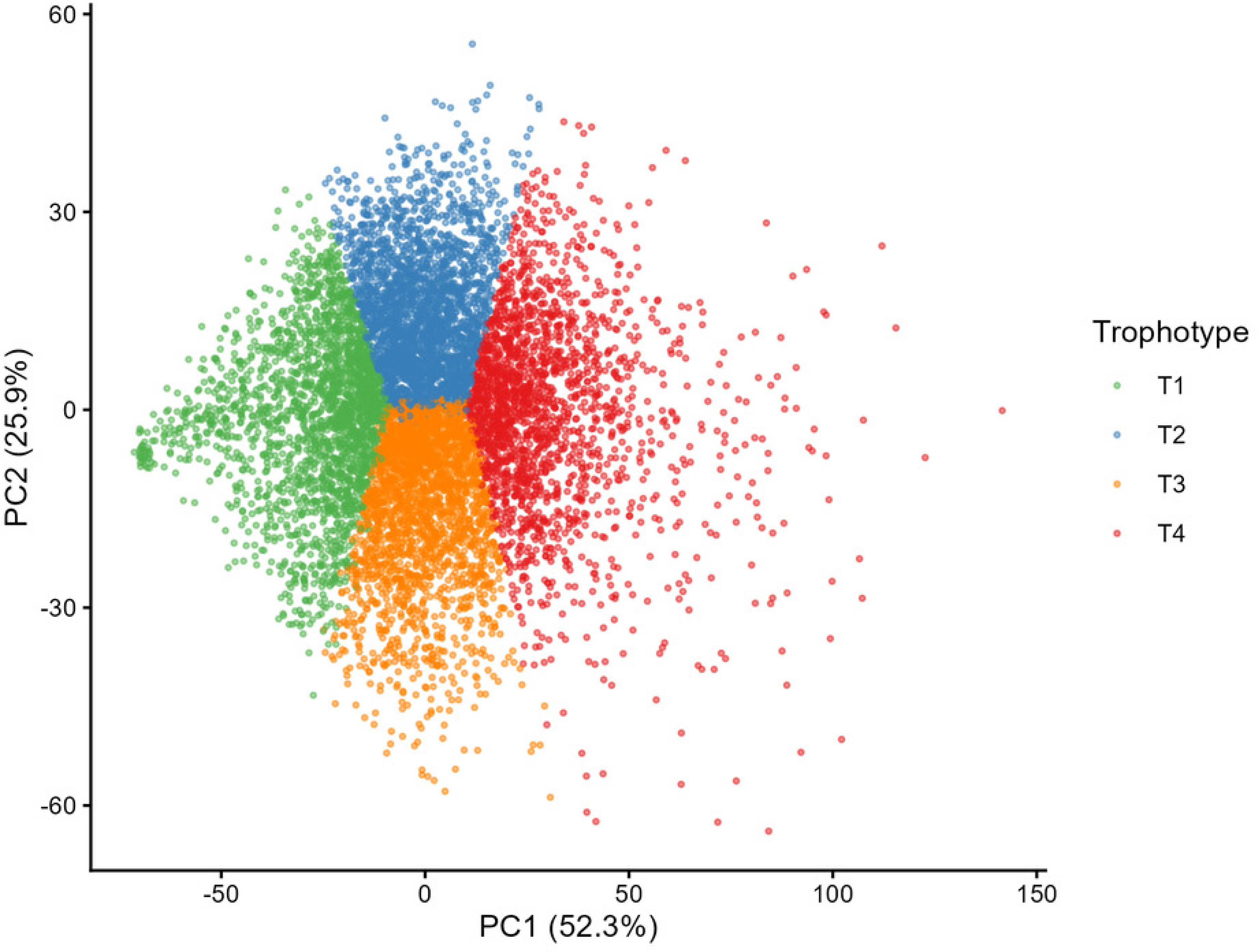
Trophotypes in the principal subspace of the guild-richness data. Projection of the 8,960 samples onto the plane defined by the first two principal components of the twelve-dimensional guild-richness matrix. PC1 (52.3% of total variance) is a positive combination of primary generalist degraders (loading 0.66), butyrate producers (0.60) and acetate producers (0.40). PC2 (25.9%) is dominated by primary generalist degraders (0.75) and butyrate producers (−0.53). Together, the first two components explain 78.2% of the total variance of the system. Each sample is coloured by its AMD-derived Trophotype: T1 (green), T2 (blue), T3 (orange), T4 (red). The four configurations form compact, contiguous regions of the principal plane, reproducing in the empirically defined subspace the four-quadrant arrangement visible in Fig. 2A. The principal subspace identified empirically coincides with the subspace defined by the guilds that thermodynamic theory of microbial communities identifies as the macroscopic axes of the colonic energy-flow network (9). Full loadings for PC1, PC2 and PC3 are reported in Supplementary Table S2.

These three guilds also map naturally onto current theory of microbial community dynamics. Primary generalist degraders open the energetic flux into the fermentative network by processing complex polysaccharides arriving in the colon; butyrate and acetate producers retain that energy through short-chain fatty acids — butyrate via classical clostridial fermentation, acetate via the Wood–Ljungdahl pathway. Marsland et al. (9) identify input through primary degradation and retention through short-chain fatty acid production as the macroscopic axes of the colonic energy-flow network. The empirical structure recovered here is consistent with this prediction: two macroscopic dimensions organise the system, with the retention dimension implemented through two biochemically distinct routes whose partial decoupling is itself a structural feature of the data.

Examination of the three pairwise planes (Fig. 2A–C) reveals a structured architecture rather than a simple two-dimensional plane. The four Trophotypes separate sharply in the input × butyrate plane (Fig. 2A); the partition is broadly recovered in the input × acetate plane, with acetate distinguishing T3 from T2 and isolating T4 (Fig. 2B); in the butyrate × acetate plane the four configurations align along a positive diagonal but the region of high acetate with low butyrate is empty (Fig. 2C) — consistent with a thermodynamic coupling between butyrogenic fermentation and acetogenesis, although the precise mechanism — whether via direct H2 transfer or via formate as electron carrier in the most prevalent gut *Blautia* species (31, 32) — remains under investigation. The third principal component (8.4% of the variance), dominated by an opposition between acetate and butyrate, captures the partial decoupling between the two retention routes.

The four Trophotypes are not simply ordered along a continuous gradient. If they were, AMD would have returned a continuum or a coarse binary partition rather than four well-separated configurations with σ-equivalent comparable to that of vertebrate communities at global scale. The discreteness detected by AMD, together with the asymmetric pattern by which butyrate and acetate are sequentially activated across configurations (Table 1), supports a reading of T4 as qualitatively distinct from T1–T3: it is the only configuration in which the acetogenic route reaches full activation, jointly with butyrate-mediated retention and high input capacity.

### Host metadata weakly predict Trophotype membership

I asked whether host-level metadata available in the AGP cohort predict membership in the four Trophotypes. After cleaning an analytic dataset of 4,260 samples with complete information across thirty-nine predictors spanning demographic, dietary, gastrointestinal, lifestyle, geographic and clinical variables, I fitted three independent classifiers: random forest, evolutionary classification tree and tuned gradient-boosted classifier.

Across the three algorithms, Cohen’s κ ranged from 0.094 to 0.131 (random forest 0.125, evolutionary tree 0.094, tuned gradient boosting 0.123) — values within the “slight agreement” range of Landis & Koch (17), well below the 0.20 threshold for “fair agreement”. The subcohort restricted to participants reporting no antibiotic exposure in the past year, intended as the closest available proxy for the system in or near its stable attractor regime, did not show improved performance (κ = 0.131).

The variables marginally informative across the three algorithms are broad descriptors of long-term lifestyle: host age (mean decrease in accuracy 0.016), types of plants consumed in the past week (0.016), and body mass index (0.009) lead the importance ranking (Fig. 5). Variables that previous literature identifies as biologically influential — antibiotic history, detailed dietary patterns, bowel-movement frequency — did not contribute beyond the chance baseline. Among the marginal exceptions, the evolutionary tree identified medically diagnosed inflammatory bowel disease as a predictor of T4 membership with an error rate substantially below the chance baseline (38.6% in a leaf of 153 samples, versus 55–70% in the remaining leaves). The direction of this association — T4 has the highest mean richness of butyrate producers in the cohort, whereas IBD is consistently associated with butyrogenic depletion in the clinical literature — is at first sight paradoxical, and is examined in the Discussion. The overall pattern is consistent with the known limitations of self-reported survey data (18, 19).

### A methodological benchmark: DMM does not detect the same structure

To assess whether the discrete structure detected by AMD depends on the specific algorithmic approach, I applied Dirichlet-multinomial mixture modelling (DMM) to the same matrix for K = 1 to K = 12 components. DMM is a probabilistic, parametric alternative to AMD: it asks whether the data can be represented as a finite mixture of Dirichlet-multinomial distributions, with optimal K identified by the elbow or absolute minimum of an information criterion. The Laplace approximation to the model evidence (Fig. 4B) decreased monotonically and smoothly across the entire range, from 234,553 at K = 1 to 222,617 at K = 12, without absolute minimum and without discernible elbow; AIC and BIC trajectories were qualitatively identical (Supplementary Fig. S1). Under standard interpretation criteria (15), DMM does not identify a finite number of components as an adequate description of these data.

**Figure 4.**
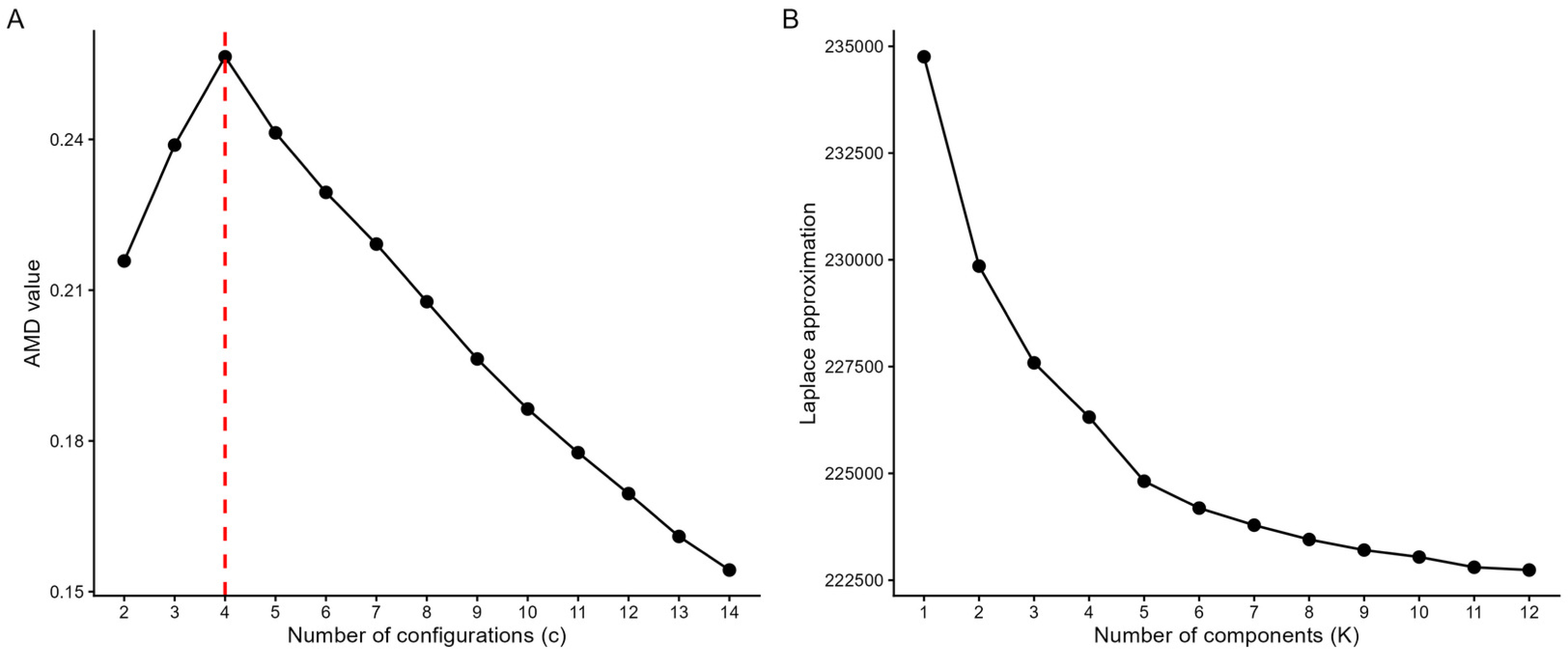
AMD detects discrete structure where DMM does not. (A) AMDi curve from Fig. 1A, reproduced for direct visual comparison with the DMM benchmark. (B) Laplace approximation to the model evidence for Dirichlet-multinomial mixture models (DMM) (15) fitted to the same guild-richness matrix for K = 1 to K = 12 mixture components. Lower values indicate better fit. The Laplace approximation decreases monotonically and smoothly across the entire tested range, from 234,553 at K = 1 to 222,617 at K = 12, without reaching an absolute minimum and without exhibiting a discernible elbow. AIC and BIC trajectories are qualitatively identical (Supplementary Fig. S1). The contrast between the two panels summarises the methodological result: AMD identifies a finite number of recurrent configurations as a geometric property of the point cloud, whereas DMM does not identify a finite number of components as an adequate description of the same data under a probabilistic-parametric model. The discrete structure detected by AMD is not reducible to a Dirichlet-multinomial mixture.

**Figure 5.**
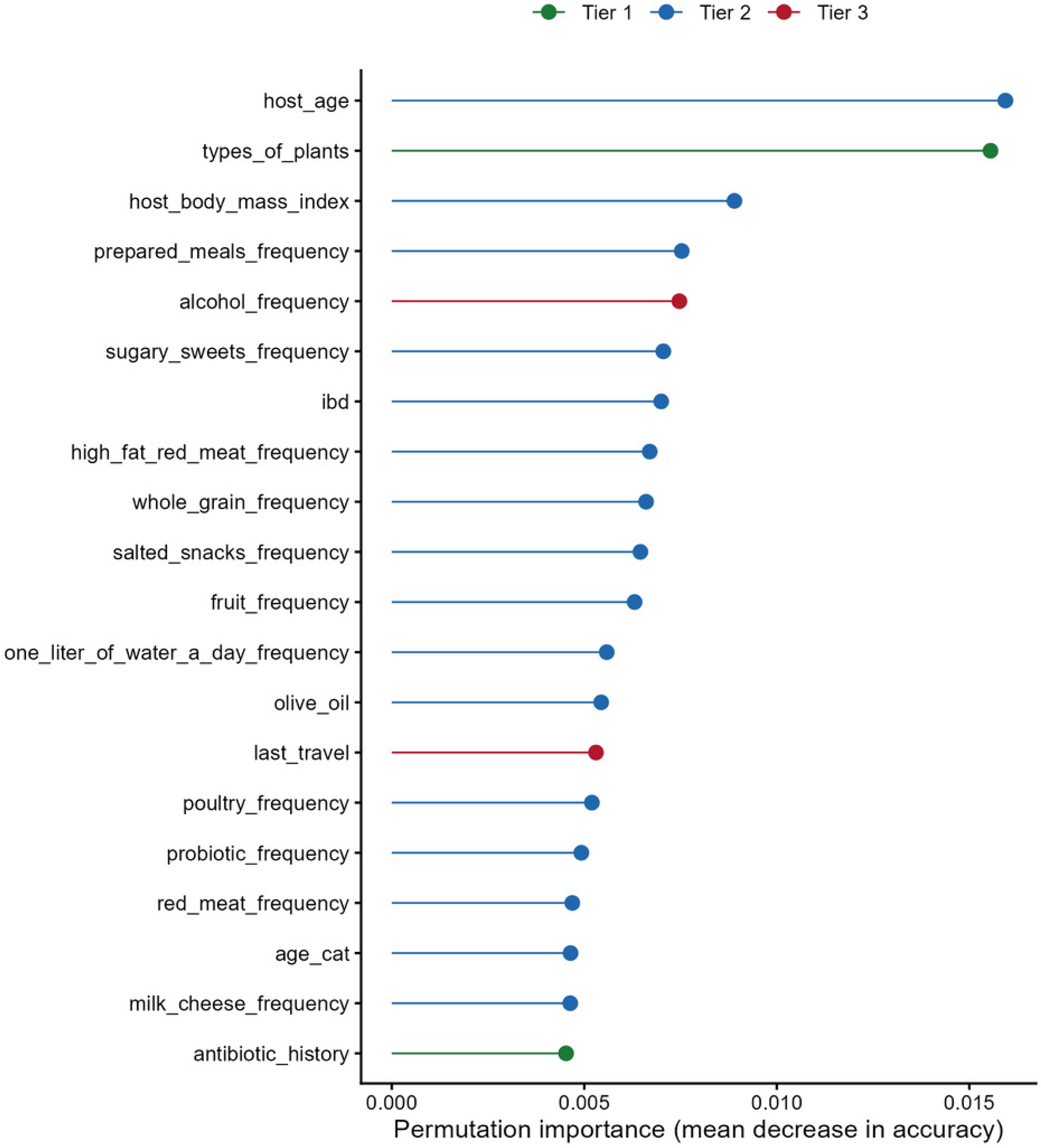
Host metadata weakly predict Trophotype membership. Variable importance from a random forest classifier predicting Trophotype assignment from thirty-nine host-level metadata variables on an analytic dataset of 4,260 samples with complete records. Importance is measured as the permutation-based mean decrease in classification accuracy on out-of-bag samples. The top twenty variables are shown, coloured by prior biological expectation: Tier 1 (green; high prior, including bowel-movement frequency and quality, antibiotic history, dietary pattern and types of plants consumed), Tier 2 (blue; moderate prior, including demographic, anthropometric, dietary-frequency, gastrointestinal and probiotic-use variables) and Tier 3 (red; weak prior, including lifestyle, geography and recent travel variables). The variables that emerge as marginally informative are broad descriptors of long-term lifestyle that integrate over many specific behaviours; variables that the literature has identified as biologically influential on microbiome composition do not contribute beyond the chance baseline. Across the three classifiers applied to this dataset, Cohen’s κ ranged from 0.094 to 0.131, values within the “slight agreement” range of Landis & Koch (17) and well below the 0.20 threshold for “fair agreement”.

The key result lies in the contrast between the two methods. AMD identifies four recurrent configurations as a clear peak (Fig. 4A); DMM returns no finite optimum on the same data. What AMD detects is a geometric structure --- regions of high local density in guild-richness space ---that is not reducible to a finite parametric mixture. This is consistent with the framing of the macroecological framework: configurations emerge as attractors of network-level dynamics, manifesting as regions of concentrated density, not as components of a probabilistic mixture over the composition space.

## Discussion

The principal result of this study is the convergence between two methodologically independent analyses on the same three functional dimensions of the human gut microbiome. AMD identifies four discrete configurations sustained by variation in three guilds — primary generalist degraders, butyrate producers and acetate producers — without any prior on which guilds should structure the system. PCA on the full guild-richness matrix, performed without reference to the AMD assignment, identifies the same three guilds as the empirical axes of greatest variation. AMD detects regions of locally elevated density and partitions the data, whereas PCA decomposes the global structure of variance without imposing any partition. Neither result follows from the other. A diagnostic random forest fitted on the AMD assignments confirms that the same three guilds dominate the partition, although this third analysis is not independent from AMD. The alignment of these three guilds with the macroscopic axes that thermodynamic theory identifies as the input and retention nodes of the colonic energy-flow network (9) supports the interpretation of the four Trophotypes as recurrent configurations of the energetic organisation of the community rather than artefacts of a particular clustering algorithm.

### Trophotypes and the literature on enterotypes

The existence of discrete clusters in the composition of the human gut microbiome has been debated since the original description of enterotypes (11). Subsequent work showed that taxonomic discreteness is less sharp than originally proposed and depends substantially on methodological choices (12); a comprehensive reassessment concluded that the composition space exhibits regions of higher density rather than hard clusters separated by empty space (13) — a geometric description that aligns with the kind of structure AMD is designed to detect. The framework adopted here suggests a reconciliation: the discreteness predicted by complex-systems theory operates on the functional organisation of the community rather than on taxonomic identity, a contingent outcome of evolutionary history and dispersal. Trophotypes and enterotypes are therefore responses to different questions formulated in different representations of the system. Taxonomic clusters, when observed, may correspond to partial projections of functional configurations onto the taxonomic axis, mediated by the loose association between taxonomy and feeding strategy in microbial systems. The discreteness emerges sharply in the functional representation because the stoichiometry of the energy-flow network constrains which combinations of input, retention and disposal of energy are viable, whereas it imposes no comparable restriction on which taxonomic identities sustain those combinations.

A complementary functional framework has been recently proposed by Meawad et al. (34), who applied non-linear archetypal analysis to 9,838 whole-genome metagenomic samples and identified three archetypes defined by metabolic pathway abundances, with most microbiomes characterised as continuous mixtures. Two methodological differences explain the discrepancy. Archetypal analysis finds the vertices of a simplex within which samples are positioned as continuous mixtures, whereas AMD detects regions of local density and returns a sharp peak only when discrete structure is present. The two also differ in the descriptor of state: pathway abundance integrates the momentary activity of the metabolic network, whereas guild richness integrates the long-term capacity to sustain it. The two frameworks are therefore complementary, and the contrast between continuous-mixture and discrete-attractor representations may itself reflect the time scale at which functional organisation is being observed.

### Why host metadata predict Trophotype membership only weakly

Across three independent classifiers, Cohen’s κ ranged from 0.094 to 0.131, within the “slight agreement” range of Landis & Koch (17). Four interpretations are compatible with this pattern. First, the relevant control parameters — host transit, dietary composition, antibiotic exposure, immune state — may be inadequately captured by self-reported survey data; objective measurements of gut transit (19) and dietary intake (18) reveal effects not recovered when assessed by participant recollection, and the only metadata variable anchored in an objective clinical procedure (medically diagnosed IBD) was indeed the only one with a predictive signal substantially below the chance baseline. Second, the relevant control parameters may be entirely absent from the cohort metadata: variables such as colonic pH, hydrogen partial pressure, mucin turnover rate or local immune cell composition are not collected at all. Third, the four Trophotypes may be robust attractors such that multiple configurations are accessible under the same host conditions; the modest κ would then reflect multistability in the sense of Dubinkina et al. (10), with empirical support for related forms of bistability reported in longitudinal genus-level data (20). Fourth, the number of Trophotypes detected here may underestimate those accessible worldwide: the AGP cohort is geographically biased toward North America, and configurations associated with diet ranges atypical of the cohort would be under-represented. The four interpretations are not mutually exclusive, and the cross-sectional design does not discriminate among them.

A specific feature of the metadata analysis deserves separate discussion. The evolutionary tree identified medically diagnosed IBD as a predictor of T4 membership with an error rate substantially below the chance baseline. The direction of this association is at first sight paradoxical: T4 has the highest mean richness of butyrate producers in the cohort, whereas the clinical literature consistently associates IBD with depletion of butyrogenic taxa. The most direct reconciliation comes from the framework of this study itself: the state descriptor is OTU richness per guild, not abundance, and the two can diverge. In IBD, multiple butyrogenic taxa may remain present at low and fluctuating abundance, yielding high observed richness despite low butyrate flux — consistent with the broader observation that disbiotic communities show diversity profiles dissociated from functional output. Additional contributing factors are difficult to rule out: AGP self-selection probably over-represents patients in remission or with milder disease; clinical heterogeneity is invisible to the AGP metadata; the leaf of 153 samples is modest. These caveats do not invalidate the finding, but they do mean that the association should be examined in cohorts with finer clinical phenotyping.

### Energy as an organising dimension

The convergence between the empirical structure of the human gut microbiome and the thermodynamic theory of microbial communities is consistent with a broader pattern in previous applications of the macroecological framework. Trophic structures of terrestrial mammal communities organise along a gradient of net primary productivity (5); the same six structures recur in birds and mammals combined despite differences in guild definitions (6); marine vertebrate communities recapitulate the pattern (7). In all of these the dimensions that structure the configurations have direct energetic interpretation. The present study extends the pattern to microbial systems with two differences. First, the configurations were detected at a much shorter time scale — a single faecal sample reflects the system over weeks to months, whereas the trophic structure of vertebrate communities results from ecological and evolutionary processes operating over substantially longer time scales. Second, the energetic dimensions identified here are state variables of the community, not external control parameters: counts of distinct organisms within metabolic guilds, not climate variables imposed from outside.

### Limitations

Six limitations should be declared. First, the design is cross-sectional: the dynamics that would confirm the attractor interpretation cannot be observed directly. Second, the AGP cohort is biased toward North America and toward adults self-selecting into a citizen-science protocol; whether the four Trophotypes recover in non-Westernised populations remains open. Third, the OTU table is built against Greengenes 13.8, superseded by SILVA 138 and GTDB; because the analysis operates at genus level via a curated dictionary, errors at finer resolution are absorbed, but a reanalysis with an updated reference would be a useful sensitivity check. Fourth, the guild dictionary covers eighteen genera; functional capacity sustained by less abundant or less characterised genera is not captured. Fifth, guild richness integrates long-term capacity at the cost of discarding information about the dominance of individual organisms. Sixth, the host metadata available in the AGP are self-reported and incomplete relative to plausible control parameters.

### Future directions

Three lines of investigation follow. First, longitudinal cohorts with serial sampling and objectively measured control parameters would allow direct tests of the multistability hypothesis: stable individuals should remain within a single Trophotype across time, transitions should be detectable, and their kinetics should inform the basin-of-attraction landscape. Second, testing the framework in cohorts with substantially different cultural and dietary backgrounds would show whether the same four configurations generalise, and might reveal additional ones associated with diet ranges not represented in the AGP. Third, the framework can be extended to other microbial ecosystems — soil, marine — to test whether the property of organising into a small number of recurrent functional configurations is generic or specific to the human colon.

## Acknowledgements

The author acknowledges the use of an AI-assisted tool for language editing and code review during manuscript preparation; all scientific content, methodological decisions and interpretations are the sole responsibility of the author.

## Author contributions

MM conceived the study, conducted all analyses, prepared the figures and wrote the manuscript.

## Data availability

The complete analysis pipeline (R scripts), the metadata cleaning specification, the guild-richness matrix, the Trophotype labels and the genus-to-guild dictionary are deposited at Zenodo (21). The raw 16S sequences and original metadata are available via Qiita (study 10317) under the terms of the American Gut Project open data agreement.

## Figure legends

**Supplementary Figure S1.**
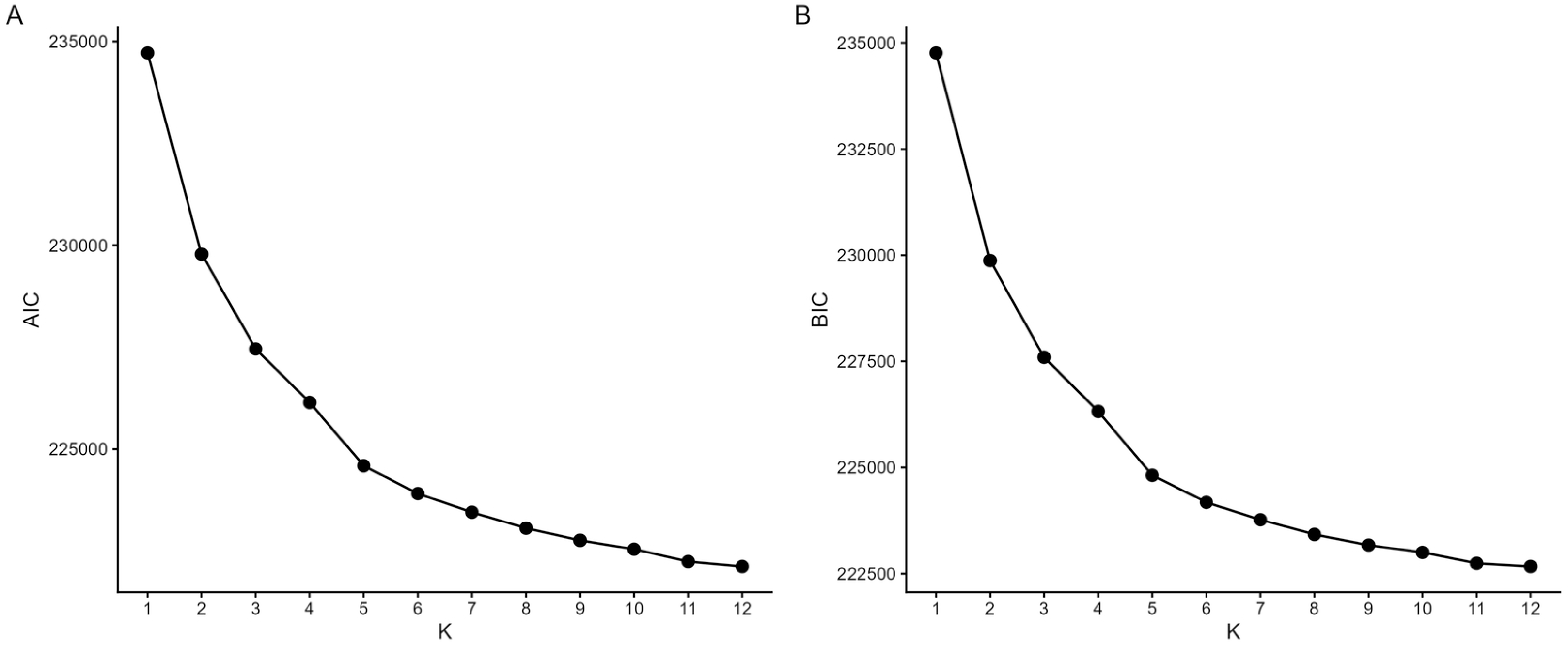
DMM benchmark — AIC and BIC trajectories. (A) Akaike information criterion (AIC) and (B) Bayesian information criterion (BIC) for the Dirichlet-multinomial mixture models fitted to the guild-richness matrix for K = 1 to K = 12 mixture components. Lower values indicate better fit. Both criteria decrease monotonically and smoothly across the tested range without reaching an absolute minimum and without exhibiting a discernible elbow, in agreement with the Laplace approximation shown in Fig. 4B.

## Supplementary Materials

**Table S2.** Variable loadings on the first three principal components of the guild-richness matrix.

| **Guild** | **PC1** | **PC2** | **PC3** |
| --- | --- | --- | --- |
| Primary_Degraders_Generalist | 0.657 | 0.750 | −0.021 |
| Butyrate_Producers | 0.603 | −0.526 | 0.536 |
| Acetate_Producers | 0.399 | −0.374 | −0.826 |
| ResistantStarch_Degraders | 0.198 | −0.092 | 0.096 |
| Propionate_Producers | 0.050 | −0.039 | −0.015 |
| Lactate_Producers | 0.045 | −0.078 | −0.068 |
| Primary_Degraders_PlantFiber | 0.034 | −0.072 | 0.122 |
| Mucin_Degraders | 0.011 | 0.007 | −0.002 |
| H<sub>2</sub>_Consumers_Asaccharolytic | 0.007 | 0.004 | 0.007 |
| H<sub>2</sub>_Consumers_SulfateReducers | 0.004 | 0.002 | 0.013 |
| Proteolytic_and_Opportunistic | 0.004 | 0.001 | −0.008 |
| H<sub>2</sub>_Consumers_Methanogens | 0.001 | −0.002 | 0.008 |
| **Standard deviation** | **22.845** | **16.087** | **9.185** |
| **Proportion of variance (%)** | **52.3** | **25.9** | **8.4** |
| **Cumulative variance (%)** | **52.3** | **78.2** | **86.6** |

Loadings of each of the twelve metabolic guilds on the first three principal components of the centred (but not scaled) guild-richness matrix (n = 8,960 samples). Guilds are ordered by absolute loading on PC1. The bottom rows summarise the spread of each component (standard deviation, proportion of variance explained, and cumulative variance). PC1 and PC2 jointly account for 78.2% of the total variance; PC3 adds 8.4%, dominated by an opposition between acetate and butyrate producers.

